# Quantitative mapping of protein-peptide affinity landscapes using spectrally encoded beads

**DOI:** 10.1101/306779

**Authors:** Huy Nguyen, Jagoree Roy, Björn Harink, Nikhil Damle, Brian Baxter, Kara Brower, Tanja Kortemme, Kurt Thorn, Martha Cyert, Polly Fordyce

## Abstract

Transient, regulated binding of globular protein domains to Short Linear Motifs (SLiMs) in disordered regions of other proteins drives cellular signaling. Mapping the energy landscapes of these interactions is essential for deciphering and therapeutically perturbing signaling networks, but is challenging due to their weak affinities. We present a powerful technology, MRBLE-pep, that simultaneously quantifies protein binding to a library of peptides directly synthesized on beads containing unique spectral codes. Using computational modeling and MRBLE-pep, we systematically probe binding of calcineurin (CN), a conserved protein phosphatase essential for the immune response and target of immunosuppressants, to the PxIxIT SLiM. We establish that flanking residues and post-translational modifications critically contribute to PxIxIT-CN affinity, and discover CN-inhibitory peptides with unprecedented affinity and therapeutic potential. The quantitative measurements provided by this approach will improve computational modeling efforts, elucidate a broad range of weak protein-SLiM interactions, and revolutionize our understanding of signaling networks.

## Introduction

*In vivo*, rapid regulation of weak, transient protein-protein interactions is essential for dynamically shaping cellular responses. Nearly 40% of these interactions are mediated by 3-10 amino acid Short Linear Motifs (SLiMs) interacting with protein globular domains (e.g. SH3, SH2, and PDZ domains) or enzymes (e.g. kinases and phosphatases)^1–3^. The human proteome is estimated to contain more than 100,000 of these SLiMs, many of which are highly regulated by post-translational modifications (PTMs) such as phosphorylation^3^. Measuring and predicting binding affinities for known and as-yet-undiscovered SLiM-binding interactions is essential for predicting signal strengths within signaling networks, understanding how these networks are perturbed by human disease, and identifying new therapeutic inhibitors^4^.

Unfortunately, the low affinities of SLiM-mediated interactions (*K_d_* values of ~1 to 250 µM) and the widespread prevalence of PTMs within them render SLiMs difficult to characterize experimentally. High-throughput approaches such as affinity purification coupled to mass spectrometry (AP-MS) and yeast two-hybrid (Y2H) assays fail to capture these weak interactions, with only ~1% of Y2H associations relying on SLiMs^2^. Proteome peptide phage display (Pro-PD), which presents peptides from all disordered proteome regions, is sensitive enough to detect low affinity interactions but does not provide binding affinities. In addition, identified peptides do not include PTMs, are biased towards the strongest binders, and include many false positives, requiring downstream validation of hundreds to thousands of candidate peptide ‘hits’^5^. Finally, a failure to observe a given peptide sequence in the bound population could result from a true lack of binding or simply poor expression or display. Parallel chemical synthesis approaches (e.g. SPOT arrays) allow testing of peptides including PTMs, but cannot provide quantitative information about affinities, require large amounts of purified protein, and lack in-line quality controls to ensure that the correct peptides were synthesized at each spot^6^.

We present a powerful technology for quantitatively profiling many SLiM-mediated protein-peptide interactions using very small amounts of material. Peptides are synthesized directly on spectrally encoded beads (MRBLEs, for Microspheres with Ratiometric Barcode Lanthanide Encoding)^7,8^ with a unique linkage between each peptide sequence and a given spectral code. MRBLE-pep libraries can then be pooled and assayed for protein binding in a single small volume before being imaged to identify the peptide sequence associated with each bead and quantify the amount of protein bound. On-MRBLE chemical synthesis allows for precise control of peptide density, incorporation of PTMs at known locations, and in-line assessment of peptide quality via mass spectrometry to identify amino acids that ablate and promote protein binding with equal confidence. In addition, MRBLEs have been engineered to have slow on- and off-rates, thereby allowing quantitative measurement of weak interactions difficult to detect via other techniques.

Here, we apply MRBLE-pep towards the study of calcineurin (CN), a conserved Ca^2+^/calmodulin-dependent phosphatase that relies on SLiMs for substrate recognition. Although CN plays critical roles in the human immune, nervous, and cardiovascular systems and likely dephosphorylates hundreds of downstream targets, only ~50 are known to date^9^. These include the NFAT family of transcription factors, whose dephosphorylation by CN is required for T-cell activation and adaptive immunity. Consequently, CN is the target of the widely used immunosuppressants cyclosporin A (CysA) and FK506. However, these drugs cause many adverse effects, including kidney failure, diabetes, pain, or seizures^10^, by inhibiting the ubiquitously expressed CN in non-immune tissues.

CN dephosphorylates sites with little sequence similarity, instead recognizing substrates by binding to two characterized SLiMs (PxIxIT and LxVP) located at variable distances from the phosphosite^11^. Blocking SLiM binding to CN prevents dephosphorylation without altering its catalytic center: FK506 and CysA prevent LxVP docking, the viral inhibitor A238L blocks PxIxIT and LxVP binding, and the high-affinity peptide inhibitor PVIVIT blocks PxIxIT binding. PxIxIT motifs vary in affinity (K s of ~1-250 µM), and determine biological output by specifying the Ca^2+^ concentration-dependence of substrate dephosphorylation in vivo^12,13^. However, the relationship between PxIxIT sequence and CN binding affinity has never been mapped systematically or probed outside of the core motif. A comprehensive understanding of PxIxIT-CN binding would allow discovery of novel CN substrates and aid efforts to rationally design CN inhibitors with enhanced selectivity and fewer side effects.

Here, we use MRBLE-pep in combination with structure-based computational modeling to systematically mutate residues at each position within and flanking three previously-characterized PxIxIT sequences and quantify their effects on affinity. We find that flanking amino acids and post-translational modifications play surprisingly critical and previously unappreciated roles in determining interaction strengths. Through iterative cycles of mutagenesis, in vitro analysis, and in vivo validation, we identify several PxIxIT peptides of unprecedented binding affinity, providing novel candidate scaffolds for the development of potent CN inhibitors. In the future, this approach can be applied to a broad range of protein-SLiM interaction to map binding energy landscapes, model signaling networks, and identify novel therapeutic inhibitors.

## Results

### MRBLE-pep experimental assay overview

We developed a new high-throughput protein-peptide interaction assay based on spectrally encoded hydrogel beads containing unique ratios of lanthanide nanophosphors (MRBLEs)^7,8^. MRBLEs containing a given code are output to a particular filtered-tip in a 96-well format or two 48-2mL reaction tubes (Fig. 1A), which is then transferred to a peptide synthesizer for functionalization and solid phase peptide synthesis (SPPS), thereby uniquely linking peptide sequences with spectral codes (Fig. 1A)^14–18^. Following SPPS, MRBLE-bound peptide libraries are pooled, incubated with an epitope-tagged protein of interest and a fluorescently-labeled antibody, washed, and imaged to reveal the peptide sequence associated with each bead (by quantifying lanthanide emissions) and the amount of protein bound (by quantifying fluorescence) (Figs. 1B,C). Unlike commercially available spectrally encoded beads (e.g. Luminex), spectral codes can be ‘read’ using a single UV excitation source, preserving the ability to multiplex binding measurements using up to 3 detection antibodies^19^.

**Figure 1.**
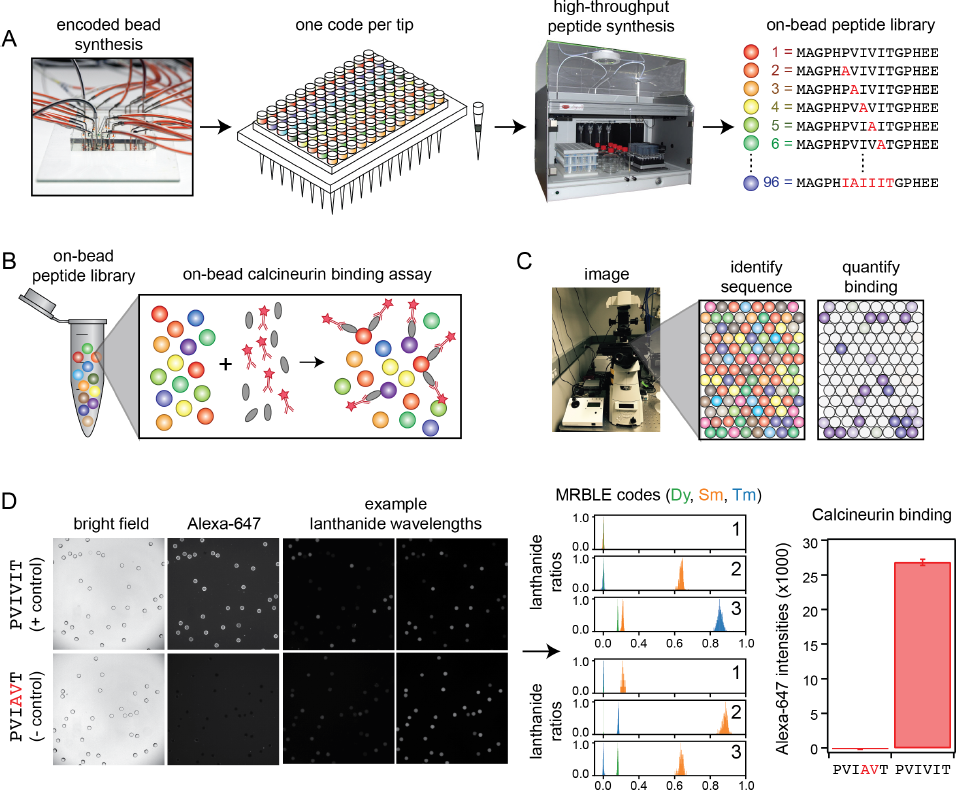
MRBLE-pep experimental pipeline for high-throughput measurement of protein-peptide interactions. (**A**) Encoded beads are synthesized in a microfluidic device and output to peptide synthesis tips arrayed in a 96 well plate format with one MRBLE code per tip. Peptides are then synthesized directly on MRBLEs via solid phase peptide synthesis with a unique 1:1 linkage between peptide sequence and spectral code. (**B**) MRBLE-pep libraries are pooled and incubated with an epitope-tagged protein and fluorescently-labeled antibody. (**C**) Following incubation and washing, peptide sequence and amount of bound protein are determined via imaging. (**D**) Example data showing images of MRBLE-pep beads coated with a CN-bound and negative control peptide, measured lanthanide ratios, and bound antibody intensities.

### Integrated synthesis quality control ensures production of full-length, correct peptide sequences

Computational prediction of novel substrates *in vivo* can be improved by including information about residues that ablate binding. However, experimentally identifying disfavored residues requires confidence that observed non-binding results from a true absence of interaction and not a failure to synthesize the correct peptide. To facilitate in-line quality assessment, MRBLEs were first functionalized with an acid-labile rink amide linker within the bead core and a non-labile glycine linker on the outer bead shell (Fig. S1)^20^; varying the ratio of extendible to non- extendible glycine linkers in the bead shell allows tuning of displayed peptide density^15,21^. Next, peptides were synthesized on both linkers via standard Fmoc SPPS. Peptides coupled to the MRBLE core via the acid-labile linker were eluted during the final acid global deprotection step and can be verified via MALDI mass spectrometry (Figs. S1, S2a-d). As peptides within the core are inaccessible to large proteins, elution allows sequence verification without reducing binding signal in downstream binding assays. Measured MRBLE lanthanide ratios before, during, and after SPPS remained constant, (Fig. S3), establishing that embedded codes are unchanged by chemical exposures.

Measuring binding affinities from a single pooled assay requires reliable estimates of the effective concentration of each peptide, which could be skewed by sequence-dependent differences in SPPS efficiency. To detect any large differences, a portion of each MRBLE-pep library was biotinylated, incubated with labeled streptavidin (DyLight650-SA), washed, and imaged to quantify bound streptavidin. Bound DyLight650-SA intensities were relatively consistent between sequences (Fig. S4) and saturated at ~20 nM DyLight650-SA for ~7100 beads in a 100 µL reaction volume (Fig. S5), establishing a surface density of ~2 × 10^8^ peptide molecules per MRBLE.

### MRBLE-pep yields quantitative measurements of calcineurin binding affinities

Measuring CN-PxIxIT binding affinities represents a demanding test for a new protein-peptide interaction assay, as CN binds PxIxIT peptide with weak affinities (Fig. 2A) via surface interactions (Fig. 2B). To determine MRBLE-pep assay sensitivity, we performed a pooled assay in which MRBLEs bearing 10 peptides (each synthesized on 3 spectral codes) were incubated with varying concentrations of His-tagged CN complexed with fluorescently-labeled anti-His antibody. These 10 peptides (‘triplicate low’) included the known NFATc1, NFATc2, AKAP79, and RCAN1 natural CN-interacting PxIxIT binding sites (Fig. 2A, Table S2)^22–25^, a set of 5 PVIVIT peptide mutants previously characterized via competitive fluorescence polarization assays^24,26^, and a scrambled negative control sequence. Consistent with previous observations, the high-affinity PVIVIT and PVIVVT variants showed strong binding, a scrambled peptide showed no binding, and the PVIAVT and PVIVIN variants showed low to intermediate binding (Fig 2C), with consistent results for the same peptide sequence synthesized on MRBLEs with different codes. NFATc2 and AKAP79 showed measurable binding and were therefore selected for further systematic mutagenesis. Although MALDI mass spectrometry confirmed successful synthesis of both RCAN1 and NFATc1 full-length peptides, binding was near the limit of detection in this assay (Fig 2C).

**Figure 2.**
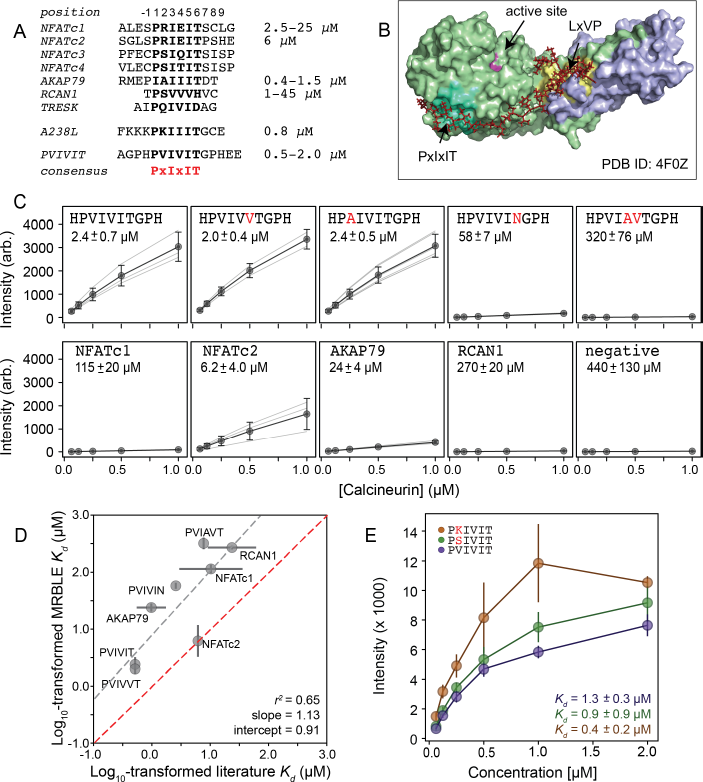
MRBLEs allow quantitative measurement of CN-SLiM interactions in high-throughput. (**A**) Sequence alignment for previously identified CN PxIxIT binding sites (see Table S1 for affinity references). (**B**) Crystal structure showing CN heterodimer bound to the A238L inhibitor including both PxIxIT and LxVP binding sites (PDB ID: 4F0Z). (**C**) Concentration-dependent binding for CN interacting with either PVIVIT variants (top), natural PxIxIT peptides (bottom), or a scrambled PxIxIT motif (bottom). Each peptide was synthesized on 3 different MRBLE codes (light grey lines); affinities were determined from global Langmuir isotherm fits (black) to median values at each concentration (grey circles). (**D**) MRBLE-derived and literature reported *K_d_* values for various CN-PxIxIT interactions. Grey solid line indicates linear regression between data sets; dotted line indicates expected 1:1 agreement. (**E**) Langmuir isotherm fits to concentration-dependent binding behavior for CN interacting with candidate high-affinity PVIVIT core variants.

Examination of known CN PxIxIT motifs (NFATc1, NFATc2) and a viral inhibitor (A238L) suggested a potential preference for positively charged (R, K) or hydroxyl (S) residues in the degenerate PxIxIT position 2 (Fig. 2A). To evaluate assay reproducibility and whether these substitutions enhance binding, we performed an additional experiment measuring binding to these same 10 peptides plus an additional 3 PVIVIT variants (PSIVIT, PRIVIT, PKIVIT) (‘triplicate high’) (Figs. 2E, Table S3). These data reproduce previously observed trends and additionally suggest that the PKIVIT peptide binds slightly more strongly than PVIVIT; therefore, we also selected PKIVIT for additional systematic mutagenesis.

To determine binding affinities for each peptide, we globally fit all data from a given assay to a single-site binding model with the assumption that while individual *K_d_* values may differ between peptides, the stoichiometry of binding remains constant (see Methods). Although these measurements take place out of equilibrium due to the need to wash beads prior to imaging, measured on- and off-rates for CN-MRBLE interactions are slow (Figs. S6, S7), suggesting that measured intensities approximate equilibrium values to yield apparent dissociation constants (Figs. S8). Slow on- and off-rates have previously been observed for hydrogel particles and are known to scale with particle radius, as proteins initially encounter and bind bead-bound ligands and then slowly diffuse into the particle via iterative dissociation and rebinding events^27^. Resultant apparent *K_d_*s largely agree with previously published values (Fig. 2D; Tables S1-3) and establish that MRBLE-pep can resolve differences in weak affinities spanning from ~0.20 to ~50 µM (a dynamic range > 2 orders of magnitude, comparable to SPR and FP^28^). However, MRBLE-pep apparent *K_d_*s slightly underestimate affinities relative to published values while preserving rank order and relative differences; therefore, we provide relative affinity differences (ΔΔG) in all subsequent analyses, which are unaffected by this shift.

### Using structure-based modeling to predict energetic effects of potential substitutions

To identify and prioritize substitutions most likely to return useful information about CN specificity, we leveraged high-resolution co-crystal structures and recently-developed computational modeling techniques^29,30^ to estimate the effects of each mutation. Co-crystal structures of CN bound to PVIVIT^24^ (PDB: 2P6B) and AKAP79^31^ (PDB: 3LL8) reveal that the conserved P1, I3, and I5 residues are buried in hydrophobic pockets on the CN surface in all three structures; for AKAP79, an I1 residue occupies the pocket typically occupied by a P1 residue. In both structures, the less-conserved solvent-exposed (positions 2 and 4) and flanking (positions −1 and 7-9) residues adopt variable side chain orientations.

We used two computational methods implemented in the protein structure prediction and design program Rosetta, the Backrub “Sequence Tolerance” protocol^29^ and a more recent “flex_ddG” method^30^. The Sequence Tolerance protocol samples rotamers of different amino acid residues at each position and returns the frequencies of amino acid residue types observed at each position in low energy sequences, thereby providing a qualitative estimate of effects of substitutions on binding. As expected, known invariant residues (P1 and I3) appeared most frequently at their respective positions; the I5 and T6 positions also tolerated I5V and T6E substitutions, respectively (Fig. 3B. These calculations additionally suggested that mutations at the variable position 2 should have little effect, but that substitution of a charged residue (E) at positions 8 and 9 could improve binding.

**Figure 3.**
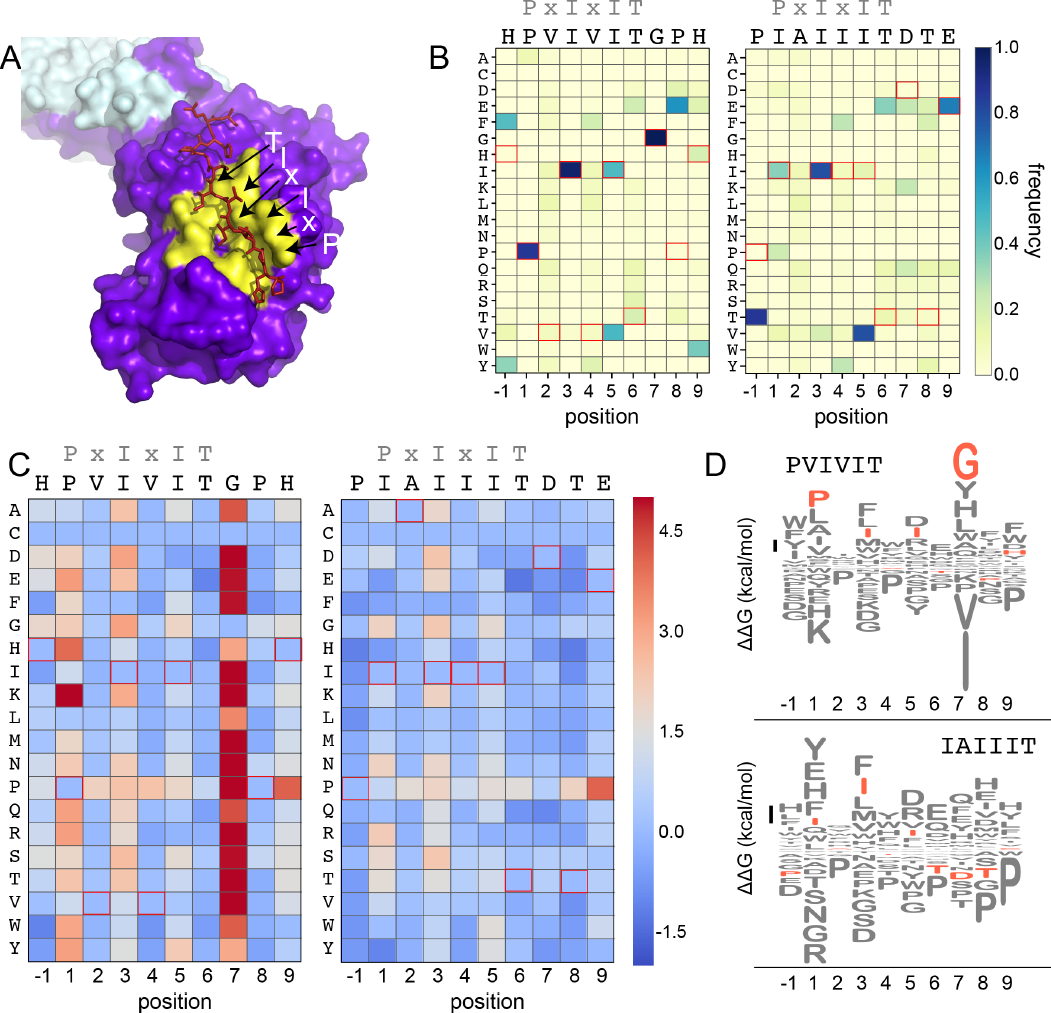
Structure-based modeling predictions. (**A**) Co-crystal structures showing zoomed in overlay of substrates. (**B**) Heat maps showing frequencies from the Rosetta Sequence Tolerance protocol for each amino acid substitution at each position for PVIVIT and IAIIIT targets. (**C**) Heat maps showing predicted changes in binding energy from the Rosetta flex_ddG protocol for each amino acid substitution at each position for PVIVIT and IAIIIT targets. (**D**) Logo motifs generated for predicted substitutions at each position within each of the 4 target motifs. Wild-type residues are shown in red; residues are arranged from top-to-bottom by predicted effect on binding energy.

While the Sequence Tolerance protocol returns ranked preferences for amino acids, it does not quantitatively predict the energetic effect of individual substitutions. To address this issue, we used the Rosetta “flex_ddG” protocol^30^ that uses protein conformational ensembles to estimate effects of mutations on binding free energies 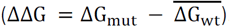. We predicted these effects for CN-PxIxIT complexes containing systematic mutations at each position (Figs. 3D, Supplementary Files). Consistent with the results of the Sequence Tolerance protocol and initial experimental results, substitutions at positions P1, I3, and I5 were generally predicted to be destabilizing (Figs. 3C,D), while changes to the solvent-exposed, degenerate x2 and x4 positions were predicted to have minimal effects (Figs. 3C,D). Intriguingly, mutations within upstream and downstream flanking residues were frequently predicted to enhance binding. Mutations to more hydrophobic or aromatic residues (W,F,Y,I,V) in the −1 position for the PVIVIT motif (Fig. 3C, left; Fig. 3D top) or to most residues other than the native P-1 for the AKAP79 motif (Fig. 3C, right; Fig. 3D bottom) improved binding *in silico*, as did mutations at Position 8 to aromatic (F,Y), aliphatic (I, V, L), or acidic (D, E) residues (Figs. 3C,D).

### High-throughput MRBLE-pep mapping of the CN-PxIxIT binding energy landscape

Using these computational predictions as a guide, we experimentally mapped the CN-PxIxIT binding energy landscape by measuring CN binding to two MRBLE-pep libraries for each of the PVIVIT, PKIVIT, NFATc2, and AKAP79 sequences containing systematic mutations in either the “core” (positions 1-6) or “flanking” PxIxIT residues (positions −1 and 7-9) (~368 peptides total) (Figs. 4A, S9-S16, Tables S12-19). The optimal number of peptides to screen per assay depends on the competing effects of ligand depletion and competition, the fraction of peptides expected to bind, the range of interaction strengths, and the statistical robustness of each measurement^32^. To reduce error and maximize the ability to resolve subtle differences, we profiled 48 peptides per reaction, with ~100 beads per sequence in each assay. To ensure that measured binding resulted from a true CN-PxIxIT interaction and not nonspecific binding, we repeated all experiments at a single, high concentration (250 nM) using a CN N330A/I1331A/R332A mutant defective in PxIxIT binding (CN NIR)^33^ and labeled antibody alone (Figs. S17-20, Tables S20-27). Several peptides were strongly bound by anti-His antibody but did not show binding when the antibody was pre-incubated with His-tagged CN; peptides strongly bound by the CN NIR mutant (*e.g.* PRIRIT) were removed from downstream analysis. All PVIVIT, PKIVIT, and NFATc2 libraries and the AKAP79 “core” mutation library were fit by a single-site binding model, permitting direct measurement of ΔΔG for each substitution. As measured intensities for the AKAP79 “flank” mutation library never reached saturation, MRBLE-pep measurements provide only qualitative estimates of effects on affinity.

To visualize the binding affinity landscape of CN-PxIxIT interactions, we generated graphs showing the relative change in affinity upon substitution to each mutant amino acid at each position (Figs. 4B, S21-29); information at all positions can be combined to produce a heat map of relative changes in binding energies (ΔΔG) (Fig. 4C) or scaled logos (Fig. S30). Consistent with computational predictions, the conserved P1, I3, and I5 residues in the core tolerate few substitutions, with nearly all mutations leading to a dramatic loss in affinity. The sole exception to this rule is the previously identified I5V mutation in both the PVIVIT and PKIVIT libraries that bound with an affinity similar to WT (Figs. 4C, S21-24)^24^. In contrast with computational predictions, substitutions at the variable x2 and x4 solvent-exposed positions strongly influenced affinity in a context-dependent manner. In particular, V2R and V2K mutations in the PVIVIT scaffold significantly enhanced binding, consistent with previous observations of high-affinity CN binding to the viral inhibitor containing a positively-charged residue at this position (PKIIIT)^22^.

**Figure 4.**
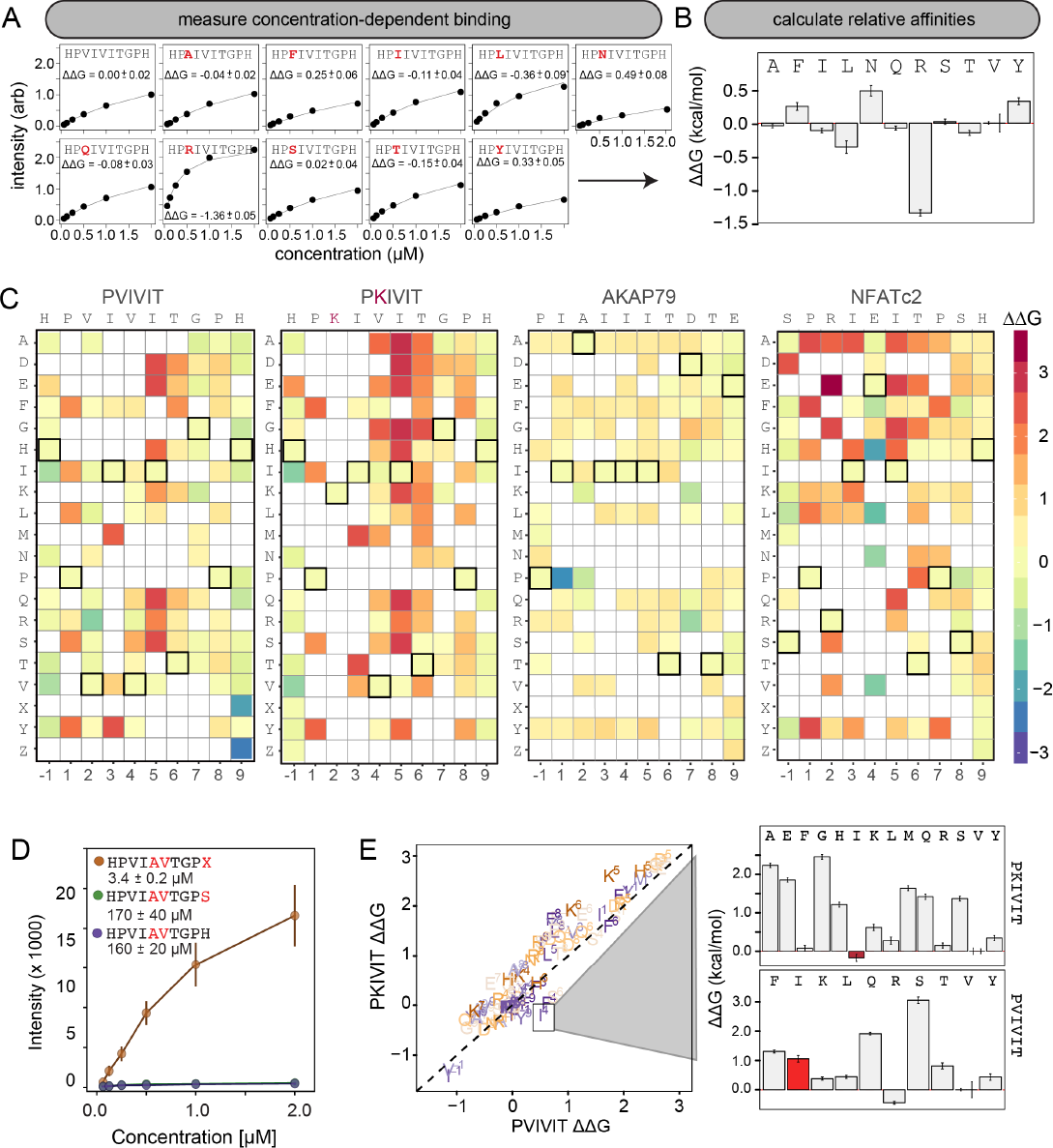
MRBLE-pep mapping of the CN-PxIxIT binding energy landscape. (**A**) Langmuir isotherm fits to concentration-dependent binding behavior for CN interacting with PVIVIT variants containing systematic mutations within position 2. (**B**) Relative change in binding affinity for individual substitutions at position 2 (as compared to WT). (**C**) Heat map showing relative affinities per substitution per position for 4 PxIxIT motifs. (**D**) Langmuir isotherm fits to concentration-dependent binding behavior for CN interacting with a medium-affinity PVIVIT variant containing the wild type histidine, a serine, and a phosphoserine at position 9. (**E**) Comparison of the effect on affinity for a particular substitution at a particular position in the PKIVIT motif (*y* axis) vs. the PVIVIT motif (*x* axis). Inset shows relative effects on affinity for various substitutions at the V4 position in PKIVIT (top) and PVIVIT (bottom), highlighting differential effects of an I substitution.

### Flanking residues play a major role in defining binding affinity and specificity

Although Rosetta modeling predicted that changes to PxIxIT ‘flanking’ residues (positions −1 and 7-9) would affect interaction affinities, the experimental impact of sequences flanking the core ‘PxIxIT’ residues on CN binding has never been probed systematically. At the −1 position, MRBLE-pep data revealed a clear CN preference for hydrophobic residues (V,L,I) and a weak preference for polar residues (T, Y, H, N, and R) for all three of the PVIVIT, PKIVIT, and NFATc2 scaffolds. By contrast, substitution to acidic (D, E) residues at this position strongly reduced binding (Figs. 4C, S9-16, S21-29). At position 9, the majority of mutations in the PVIVIT scaffold increased affinity. Substitution of phosphomimetic residues (D, E) or unphosphorylated serine increased affinity slightly and not at all, respectively, but substitution of phosphoserine and phosphothreonine residues (X, Z) increased affinity nearly 50-fold (from *K_d_* = 170 +/- 40 nM to *K_d_* = 3.4 +/- 0.2 nM) (Figs. 4C-D, S10). This effect was specific to the PVIVIT sequence, with significantly less drastic increases and decreases in affinity observed for the same substitutions in the NFATc2 and AKAP79 scaffolds, respectively (Fig. S14, S16, S31). These MRBLE-pep data establish that flanking residues make major contributions to affinity and the PxIxIT SLiM is significantly longer than previously thought.

### Effects of individual substitutions show evidence of non-additivity

Understanding the degree to which CN-SLiM binding specificity can be explained by a linear additive model is critical for estimating the likely accuracy of downstream efforts to identify substrates or design therapeutics. Deviations from additivity can be quantified via double-mutant cycle (DMC) analysis, in which the effects of individual mutations are measured alone and in combination and then compared. DMC analysis between the PVIVIT and PKIVIT scaffolds revealed that the I3Y substitution resulted in a significant loss of affinity within the PVIVIT context but had only minor effects on PKIVIT binding; by contrast, V4F and V4I mutations of this solvent-exposed position significantly increased binding only within the PKIVIT sequence (Fig. 4E). This last substitution suggests that the high affinity of the PKIIIT sequence in the A238L inhibitor in particular relies on cooperativity between the position 2 and 3 residues.

### Computational methods have limited predictive power for solvent-exposed and flexible residues

To determine the degree to which Rosetta modeling successfully predicts observed binding, we generated scatter plots showing relationships between measured affinities and predicted frequencies or changes in free energy of binding (ΔΔG), respectively, for each residue at each position (Figs. 5A, S32-33). Both the Sequence Tolerance and the flex_ddG methods correctly predicted that I3 and I5 mutations decrease affinity, and Rosetta correctly predicted that substitution to a hydrophobic I or aromatic Y residue at the −1 position would improve binding while substitution to the acidic residue E would reduce binding. However, computational modeling predicted that variations at solvent-exposed positions (V2 and V4) would have little effect on affinity, in direct contrast to experimental observations (Figs. 5A, right panels, S32-33). These results highlight the difficulties associated with modeling binding contributions for substitutions at solvent-exposed as well as flanking positions (shown in S32-S33), where effects on conformations in the bound and unbound state ensembles are unlikely to be correctly modeled.

To derive a quantitative metric for the overall predictive power of Rosetta modeling, we employed receiver-operator characteristic (ROC) curve analysis. For the predictions of the Sequence Tolerance protocol, PVIVIT variants were classified as ‘bound’ or ‘unbound’ according to whether experimentally measured *K_d_*s were 4-fold lower than the *K_d_* measured for a ‘scrambled’ peptide sequence; for the flex_ddG predictions, mutations were classified according to whether they enhanced or decreased binding relative to wild-type, yielding negative or positive ΔΔG values (Fig. 5B, C). In both cases, *in silico* predictions based on actual paired measurements modestly outperformed predictions based on shuffled data, with area under the curve (AUC) values between 0.6 and 0.7.

**Figure 5.**
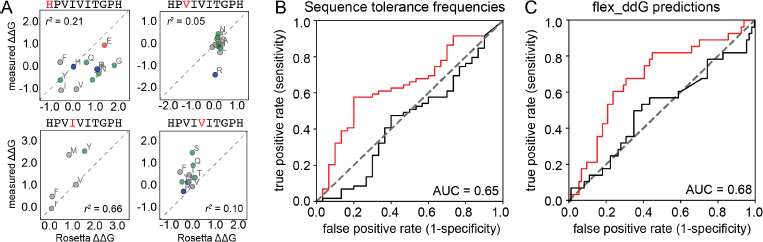
Comparisons between *in silico* modeling and experimental results. (**A**) Scatter plots showing the relationship between the experimentally measured (*y* axis) and computationally predicted (*x* axis) ΔΔG values for all experimentally tested residues within the PVIVIT sequence at Positions −1 and 3 (left) and 2 and 4 (right). (**B**) ROC curve showing calculated true positive rate (TPR, *y* axis) plotted against false positive rate (FPR, *x* axis) for classifying either real (red) or shuffled (black) data as ‘bound’ or ‘unbound’ at varying frequency thresholds. (**C**) ROC curve showing calculated true positive rate (TPR, *y* axis) plotted against false positive rate (FPR, *x* axis) for correctly predicting whether substitutions enhance or reduce affinities at varying ΔΔG thresholds computed by the Rosetta flex_ddG protocol.

### Absolute binding affinities confirm that mutations to flanking residues dramatically change affinities

To determine absolute affinities for peptides spanning a range of binding behaviors, we performed 2 experiments measuring concentration-dependent CN binding to an additional ‘calibration’ library containing ~6 peptides from each core and flank library (Figs. S34-S35, Tables S4-S5). These experiments represented full technical replicates in which peptides were synthesized *de novo* on MRBLEs produced at different times and assayed for binding using different batches of purified calcineurin. Measured ΔΔG values relative to the PVIVIT variant between experiments showed strong agreement (Pearson *r^2^* = 0.72 (Fig. S36)), demonstrating the robustness of the assay (Fig. S37). Averaged absolute affinities estimated for all variants using these ΔΔG values and the known literature *K_d_* for PVIVIT confirm that hydrophobic residues at position −1 and phosphorylated residues at position 9 significantly increase affinity by 10-fold and 100-fold, respectively (Fig. S37-S39).

### Validation of in vitro results using in vivo calcineurin activity assays

The ultimate goal of mapping CN-SLiM binding energy landscapes is to improve understanding of CN target substrate recognition and enable rational design of *in vivo* inhibitors. To test the degree to which *in vitro* MRBLE-pep affinities predict *in vivo* inhibition, we used a previously developed dual luciferase reporter assay to assess the activity of the NFAT2 transcription factor in HEK293T cells (Fig. 6A)^22^. NFAT2 must be dephosphorylated by CN to accumulate in the nucleus, where it activates transcription of its target genes, and this interaction can be inhibited by peptides or small molecules that binds to CN. Reduction of NFAT2-dependent transcriptional activity upon expression of a competing PxIxIT peptide therefore reflects, quantitatively, the affinity of the inhibitor peptide for CN. Candidate PxIxIT inhibitors were co-expressed as GFP-fusion proteins within cells to enhance their stability and facilitate direct measurement of inhibitor concentrations.

**Figure 6.**
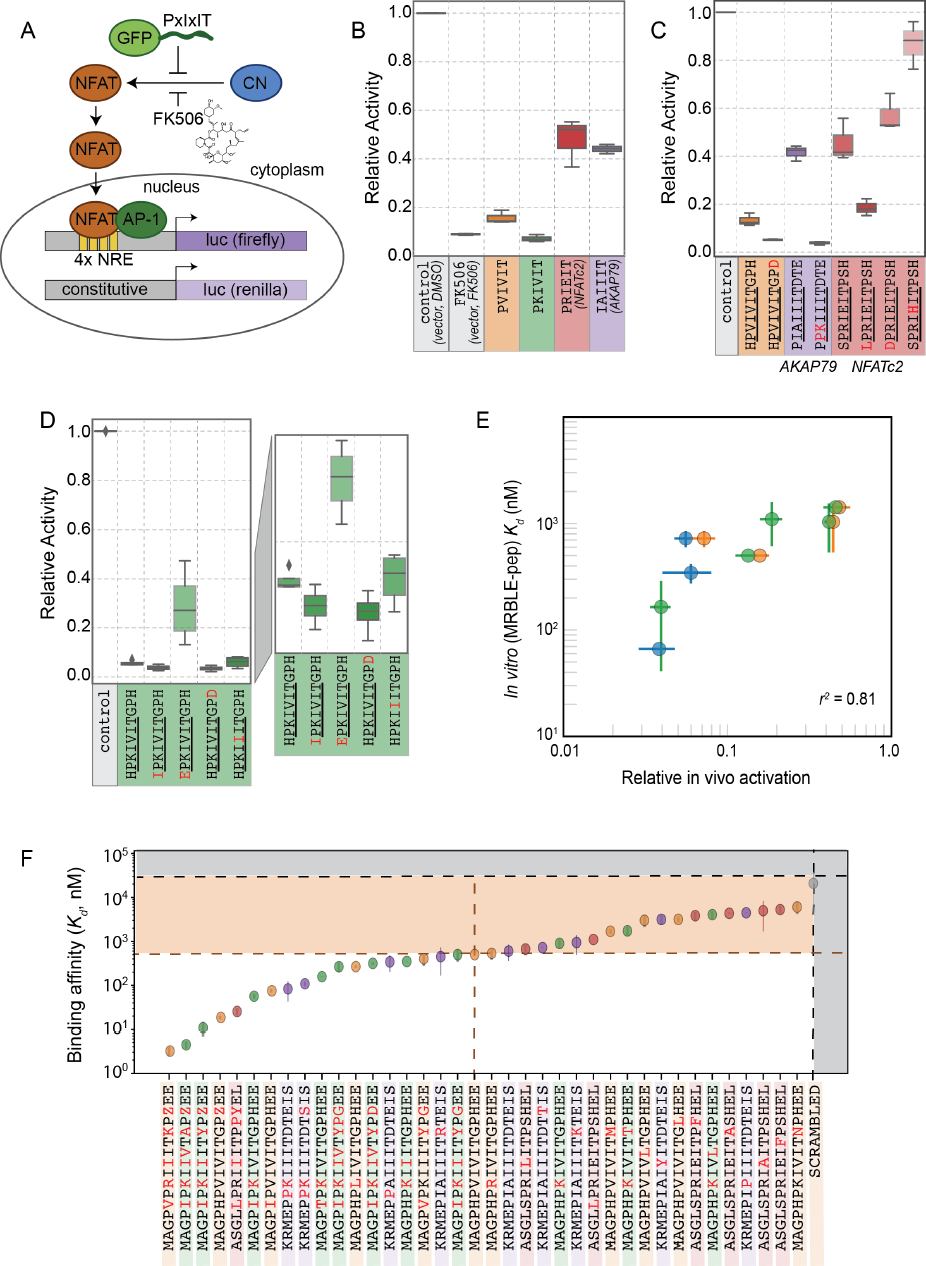
*In vivo* assays to quantify CN inhibition and an *in vitro* search for a potent inhibitor. (**A**) *In vivo* experimental assay schematic. (**B**) Reporter gene inhibition in the presence of an empty eGFP vector (control), an empty eGFP vector with topical application of FK506, and vectors expressing PVIVIT, PKIVIT, NFATc2, and AKAP79 PxIxIT peptides with a C-terminal eGFP tag. (**C**) Reporter gene inhibition in the presence of various PVIVIT-, AKAP79-, and NFATc2-eGFP variants. (**D**) Reporter gene inhibition in the presence of various PKIVIT-eGFP variants. (**E**) *In vitro* calibrated binding affinities (*K_d_*, nM) plotted against relative *in vivo* activation for data shown in panels B (orange), C (green), and D (blue). (**F**) *In vitro* measurements of binding affinities for candidate high-affinity binders identified by combining favorable residues.

We compared levels of NFAT-driven luciferase activity in the presence of empty vector, vectors driving expression of the 4 PxIxIT peptide scaffolds probed extensively *in vitro* (PVIVIT, PKIVIT, NFATc2, and AKAP79), or FK506, a small-molecule inhibitor of CN with a previously measured IC_50_ of ~0.5 nM (Fig. 6B)^34^. All 4 PxIxIT peptides substantially inhibited NFAT-driven luciferase activity, confirming competitive inhibition of CN-NFAT-(PxIxIT) binding *in vivo*. Consistent with MRBLE-pep measurements, PKIVIT was the most effective inhibitor, followed closely by PVIVIT. NFATc2 and AKAP79 showed weaker *in vivo* inhibition, consistent with MRBLE-pep results and in contrast with prior work suggesting that AKAP79 binds with equal affinity to PVIVIT (*K_d_* ~500 nM)^31^. Co-expression of the PKIVIT peptide resulted in greater inhibition than FK506, which interferes with CN-substrate interactions by occluding the LxVP-SLiM binding site^35^ and is a potent CN inhibitor.

We next examined the importance of PxIxIT flanking residues by creating substitutions in the −1 and 9 positions (Fig. 6C,D). Consistent with *in silico* predictions and MRBLE-pep affinities, substitution of a hydrophobic residue (IPKIVIT or LPRIEIT) at the −1 position significantly increased inhibition, whereas incorporating an acidic residue (EPKIVIT or DPRIEIT) greatly diminished inhibition. As predicted computationally and observed in the MRBLE-pep assays, introduction of the phosphomimetic mutation H9D resulted in slightly increased inhibition; it was not possible to directly test the extremely high affinity phosphoserine or phosphothreonine *in vivo*. Taken together, *in silico* modeling, the MRBLE-pep affinities, and *in vivo* results suggest that phosphorylation at position 9 may be a key determinant of modulating PxIxIT-CN affinity.

Finally, we sought to test whether MRBLE-pep *in vitro* affinity measurements could identify initial peptide scaffolds exhibiting strong inhibition *in vivo* for future optimization as peptide or small-molecule inhibitors. PKIVIT was a significantly stronger *in vivo* inhibitor than the previously patented PVIVIT peptide^36^, with activity comparable to the PKIIIT site within the known A238L inhibitor. As predicted, both the IPKIVITGPH and HPKIVITGPD variants exhibited even stronger binding and inhibition (Fig. 6D); similarly, mutating the I1 and A2 residues in the AKAP79 PIAIIIT motif to create a novel PxIxIT sites with adjacent prolines and a positively charged residue in the second position (PPKIIIT) greatly enhanced inhibition. Overall, MRBLE-pep *in vitro* measurements were strongly predictive of *in vivo* inhibition (Fig. 6E)

### Identification of high-affinity scaffolds with therapeutic potential

Although cyclosporin A and FK506 are routinely prescribed to transplant patients to inhibit CN-dependent immune response activation, both drugs are associated with adverse effects that likely result from inhibition of CN-substrate dephosphorylation in non-immune tissues. High-affinity peptides that inhibit a particular subset of CN-SLiM interactions could therefore serve as initial scaffolds for medicinal chemistry efforts to identify inhibitors with improved specificity. To identify candidate with high-affinity variants, we measured CN binding to a final MRBLE-pep library containing the set of 36 ‘calibration’ peptides along with 11 peptides containing combinations of mutations previously shown to increase affinity (Figs. S38- S39). Measured ΔΔGs for these full technical replicates again showed remarkable agreement (*r^2^* = 0.83) (Fig. S40). PVIVIT variants combining a phosphothreonine (Z) at position 9 with hydrophobic residues (I, V) at position −1 showed the strongest binding, with measured *K_d_* values of ~ 10 nM (50x stronger than the measured PVIVIT affinity), and multiple PVIVIT, PKIVIT, and AKAP79 variants also showed enhanced binding (Figs. 6E, S39). Together, these *in vitro* and *in vivo* measurements demonstrate that systematic determination of CN-SliM binding yields peptides with a continuous range of affinities for testing in therapeutic applications.

## Discussion

Here, we demonstrate a novel strategy to quantitatively profile the binding specificity landscape for a peptide-protein interaction, thereby generating key insights required to map, model and therapeutically perturb essential signaling networks in healthy and diseased cells. Compared to existing technologies, our approach has several advantages. Although yeast and phage display can powerfully discriminate between ‘bound’ and ‘unbound’ peptide populations^37,38^ for > 10^8^ protein-peptide interactions, these high-throughput screening methods cannot quantitatively measure affinities and are unable to probe the effects of PTMs or unnatural amino acids with therapeutic potential. The MRBLE-pep assay is also faster and requires less in material than alternative methods: measuring interaction affinities for 384 peptides using MRBLE-pep requires ~400x, ~700x, and ~9600x less purified protein than surface plasmon resonance, fluorescence polarization, and isothermal calorimetry, respectively, with savings increasing with library size (Table S6). This reduction in material should allow future profiling of a wide range of biologically important SLiM-binding domains, including those that are unstable and/or difficult to express recombinantly. MRBLE-pep libraries can be synthesized and imaged over the course of days, facilitating iterative rounds of synthesis and measurement for efficient landscape mapping. Finally, the results presented here highlight the limitations of existing *in silico* modeling methods for accurately predicting peptide-protein interactions, particularly for solvent-exposed and flexible residues. By iteratively generating and experimentally testing quantitative computational predictions of how amino acid sequence affects SLiM affinity, analyses such as these can simultaneously map a specific protein-SLiM interaction landscape and provide critical experimental data for use in revising computational algorithms. Furthermore, this approach can be extended to discover binding specificities of as-yet-uncharacterized protein domains.

Our findings reveal biologically significant insights into CN-PxIxIT specificity and illustrate the power of systematic analyses to shed light on elusive SLiM-protein interactions. The *in silico* modeling, MRBLE-pep data, and *in vivo* validation presented here are the first to systematically capture the impact of PxIxIT flanking residues on affinity, allowing identification of several peptides whose affinity for CN is two orders of magnitude higher than that of PVIVIT that are more potent inhibitors of CN-NFAT signaling *in vivo*. Furthermore, we identify a collection of peptides with a continuous spectrum of affinities for CN, which may allow inhibition to be fine-tuned *in vivo*. Currently, doctors monitor the dosage of CN inhibitors to minimize adverse consequences of these drugs. Instead, a peptide that selectively perturbs CN binding to a subset of its targets could maximize therapeutic effects, while minimizing the disruption of CN signaling events that promote health.

In addition to identifying candidates for therapeutic manipulation of CN signaling, the demonstration that residues both upstream and downstream contribute to PxIxIT-CN affinity expands the definition of this motif. Current computational strategies attempt to identify novel CN substrates by searching for sequences within the proteome that match a consensus expression for the six core residues alone. The expanded 10-residue definition of the PxIxIT motif derived here will enhance future computational efforts to comprehensively map the calcineurin signaling network, reducing the number of both false positive and false negative substrates. In particular, MRBLE-pep provides critical Information about residues that reduce affinity (*e.g.* acidic residues at the −1 position reduces affinity) that is never revealed by positive screening methods and is rarely collected due to the resource-intensive nature of such analyses using established techniques. Finally, the ability to predict affinity based on SLiM sequence is an essential step toward accurate modeling of signaling dynamics *in vivo*, as the extent of *in vivo* dephosphorylation has previously been shown to depend on primary PxIxIT amino acid sequence (and the associated interaction affinity)^12^. Even subtle (~2-fold) differences in binding could have profound effects on downstream signaling under physiologic conditions given the low affinities of CN-PxIxIT interaction.

Establishing the impact of post translational modifications on SLiM-protein affinity is critical for understanding cross talk between signaling networks *in vivo*. Consistent with our findings that acidic residues at the −1 position decrease affinity, JNK kinase regulates CN signaling by phosphorylating a serine that immediately precedes the PxIxIT of NFAT4^39^. Similarly, our demonstration that phosphorylated residues downstream of the core PxIxIT sequence at position 9 enhance PVIVIT affinity echoes observations that phosphorylation of at threonine in this position is required for binding of CN to a PxIxIT site in C16ORF74, and for its ability to promote invasiveness of pancreatic ductal adenocarcinoma (PDAC)^40^. This positive effect on binding is context-dependent, increasing PVIVIT binding 50-fold but having little or no effect on other PxIxIT sequences, reinforcing the importance of systematic analyses for generating predictive information.

Beyond improving the ability to reconstruct downstream CN signaling networks *in vivo* and identifying candidate therapeutic inhibitors, these results also have direct relevance to precision medicine. Current annotations of 2RQ, 7PA, and 7PL missense mutations within the NFATc2 PxIxIT site recovered from sequenced lung and breast adenocarcinomas describe them as unlikely to have functional effects *in vivo*^41^. While the experiments presented here did not directly test the functional consequences of a R2Q mutation, all mutations away from the native residue at this position resulted in a dramatic loss of binding (Fig. S25), and the P7A substitution led to a nearly complete loss of binding (Figs. S13, S25-26). Thus, binding specificity maps like those obtained here could both help clinicians identify functionally significant missense mutations, and refine the computational algorithms that predict the impact of such alleles. Overall, the approaches outlined here will help decipher the language of SLiM-domain binding events, and of the cellular signaling networks they define in both healthy and diseased cells.

## Methods

Reagents for peptide synthesis were purchased and used without further purification from NovaBiochem, AnaSpec (Fremont, CA), and Sigma-Aldrich (St. Louis, MO). All other solvents and chemical reagents were purchased from Sigma-Aldrich.

### MRBLE synthesis and collection

MRBLEs were synthesized using a previously published microfluidic device^7,8,19^ with each code collected into wells of a 96-well plate using an open-source in-house fraction collector. Briefly, all lanthanide input mixtures contained double-distilled water (ddH_2_O), 42.8% v/v 700 MW PEG-diacrylate (PEG-DA) (Sigma-Aldrich), 19.6 mg mL-1 lithium phenyl-2,4,6-trimethylbenzoylphosphinate (LAP), and 5.0% v/v YVO4:Eu (at 25 mg mL-1). Individual input mixtures contained 16.3% v/v of either YVO4:Sm (25 mg mL-1), YVO4:Dy (25 mg mL-1), or YVO4:Tm (12.5 mg mL-1). Lanthanide solutions were mixed using a herringbone mixer channel and forced into droplets using a T-junction with a continuous stream of HFE7500 (3M Novec) containing 2% w/w modified ionic Krytox™ 157FSH (Miller Stephenson, Danbury, CT)^42^. Droplets were then photopolymerized with UV light from a full-spectrum 200W Xenon arc lamp (Dymax, Torrington, CT, USA). With this setup, ~3,000 beads containing each spectral code were synthesized in 70 seconds (as described previously^7,8^); for the 48-plex MRBLE library used here, each code was produced 10 times to yield ~30,000 beads per code.

### Bead functionalization (PAP protocol)

MRBLE hydrogel beads were functionalized with terminal amine handles via Michael addition by reacting available acrylates in the MRBLEs with a solution of cysteamine (50 eq) containing pyridine (50 eq) in H_2_O:DMF (1:3) for 18 hours at ambient temperature. Next, to selectively functionalize and segregate MRBLE to distinct shell regions, MRBLEs were swelled in water overnight and drained using a manifold before the addition of a solution containing Fmoc-N-hydroxysuccinimide (0.2 eq) and diisopropylethylamine (DIPEA) (0.8 eq) in dichloromethane:diethyl ether (55:45) with vigorous shaking (1600 rpm for 15 seconds followed by 30 seconds at rest) for 30 minutes^15^. To selectively functionalize MRBLE inner core regions with an acid-labile rink amide linker, MRBLEs were then treated with 4-[(2,4-Dimethoxyphenyl)(Fmoc-amino)methyl]phenoxyacetic acid (rink amide) (5.0 eq), N,N′- Diisopropylcarbodiimide (DIC) (5.0 eq), and DIPEA (10 eq) in dimethylformamide (DMF) for 1 hour and repeated twice. After removal of the Fmoc protecting group using 20% 4-methylpiperidine (4-MP) in DMF for 20 minutes, the effective on-bead peptide concentration was reduced by reacting the bead with a mixture of Fmoc-glycine-OH:Ac- N-glycine-OH (1:9) (5.0 eq), DIC(5.0 eq), and DIPEA (10 eq) for 14 hours. Following Fmoc deprotection, MRBLEs were transferred to an automated peptide synthesizer for solid phase peptide synthesis (SPPS).

### On-bead peptide synthesis

Peptide synthesis was performed using a Biotage™ Syro II automated peptide synthesizer following instructions from the manufacturer (Biotage, Charlotte, NC). During coupling steps, Fmoc-protected amino acids (10 eq) were activated with HCTU (9.8 eq) and NMM (20 eq) with coupling times of 8 minutes for standard amino acids and 25 minutes for phosphorylated amino acids. Each coupling round comprised of 2 sequential coupling reactions with the addition of fresh amino acid and coupling reagents. Deprotection was performed initially with 40% 4-MP for 2 minutes and then followed with 20% 4-MP for an additional 6 minutes. MRBLEs were then washed thoroughly with 6 rounds of DMF (~0.4mL) before the next coupling step. Before global deprotection with TFA, ~50 µL from a volume of 400 µL for each code was saved for biotin conjugation (described below).

To perform the global deprotection step, MRBLEs were washed with DCM and dried under vacuum before the addition of 0.5 mL of TFA cocktail (Reagent B, TFA:phenol:ddH_2_O:triisopropylsilane, 88:5:5:2, v/m/v/v) was added to each reaction tube and reacted with shaking (15 seconds shaking and 1 minute rest) for 1.5 hours. After TFA deprotection, MRBLEs containing deprotected peptides were washed with TFA (~0.5 mL, and collected for MALDI analysis), DCM (~2 mL), neutralized with 10% DIPEA in DMF twice (~1 mL), and then finally washed with storage buffer (1 mL, 0.1% TBST with 0.02% NaN_3_) 3 times^43^. To assess peptide quality, the TFA solution containing cleaved/unprotected peptide was transferred to 15 mL falcon tubes using the provided liquid transfer system from Biotage. Peptides were triturated with cold diethylether (~1 mL), pelleted, decanted, and repeated these steps 3 times before preparing for MALDI analysis using a general protocol described below.

### Peptide concentration estimation via biotin conjugation assay

A small aliquot (~50 μL from 400 μL) of each code was transferred and pooled into a separate fritted reaction tube (Biotage 2 mL reaction tubes for SPPS) for biotin (50 eq) conjugation using DIC (50 eq), DIPEA (100 eq), and DMF overnight. After washing the MRBLEs (DMF, MeOH, and DCM), another round of coupling using the same conditions mentioned above was mixed for 2 hours and then washed. MRBLEs with terminal biotin were then globally deprotected using a cocktail of TFA/phenol/H_2_O/TIPS (87.5:5:5:2.5 v/v) at ambient temperature for 1.5 hours. After global deprotection, MRBLEs were neutralized with 10% DIPEA in DMF, washed with DCM, washed with 3 × PBS containing 0.1% TWEEN 20 (0.1% PBST), and stored in 0.1% PBST containing 0.02% NaN_3_.

After biotin conjugation, a 40 μL aliquot from a 600 μL suspension of beads was passivated with 0.1% PBST containing 5% BSA in a 150 μL PCR strip tube on a rotator overnight at ~5 °C. The beads were then washed with 0.1% PBST containing 2% BSA 3 times (~100 μL), followed by incubation with 1 μL of labeled streptavidin (final concentration = ~189 nM, Abcam, ab134241) in 0.1% PBST containing 2% BSA (99 μL) for 30 minutes on a rotator at ambient temperature. After incubation, MRBLEs were pelleted and washed with 0.1% PBST once and then imaged to obtain binding data.

### MALDI quality control

After global deprotection, supernatants collected in 15 mL Falcon tubes for each code were placed into a freezer for 1 hour; these tubes were then centrifuged (4000 g for 20 minutes at 4 °C) and decanted (repeated 3 times). Peptides were then dissolved with 60% ACN/H_2_O (20 μL, phosphopeptides were dissolved with 50% acetic acid) for MALDI analysis using THAP (250 mM in ACN) as the matrix. To prepare the MALDI plate (microScout Target MSP 96 target polished steel BC, part #8280800), 0.5 μL of sodium citrate (250 mM in H_2_O containing 0.1% TFA) was spotted on to the plate surface and allowed to dry. After drying, 1 μL of a 1:1 mixture of the peptide solution with a solution of sodium citrate:THAP (1:1) was spotted onto the plate and allowed to dry again before analysis. Data was obtained using a Bruker microflex MALDI-TOF (Billerica, MA, USA). The instrument was run on positive-ion reflector mode with a laser setting of 1,810 V and data averaged over 100 scans. Raw data was analyzed using FlexAnalysis and mMass (ver. 5.5, www.mmass.org).

### Purification of calcineurin

N-terminally, 6-His-tagged human calcineurin A isoforms (truncated at residue 400), either wild-type or containing the mutation ^330^NIR^333^-AAA, were expressed in tandem with the calcineurin B subunit in E.coli BL21 (DE3) cells (Invitrogen) and cultured in LB medium containing carbenicillin (50 mg/ml) at 37°C to mid-log phase. Expression was induced with 1 mM isopropyl 1-thio-b–galactosidase at 16°C for 18 hours. Cells were pelleted, washed and frozen at −80°C for at least 12 hours. Thawed cell pellets were re- suspended in lysis buffer (50 mM Tris-HCl pH 7.5, 150 mM NaCl, 0.1% Tween 20, 1mM β-mercapto ethanol, protease inhibitors) and lysed by sonication using four 1 minute pulses at 40% output. Extracts were clarified using two rounds of centrifugation (20,000 × g, 20 min) and then bound to 1 ml of Ni-NTA beads in lysis buffer containing 5 mM imidazole. Beads were washed with lysis buffer containing 20 mM imidazole and eluted with lysis buffer containing 300 mM imidazole. Purified calcineurin heterodimer were dialyzed in buffer (50 mM Tris-HCl pH 7.5, 150 mM NaCl, 1 mM β-mercapto ethanol) and stored in 10-15% glycerol at −80°C.

### Calcineurin binding assays (time series and dilution series)

Pre-incubation of CN with anti-His antibody significantly reduced observed background binding to sequences containing multiple basic residues due to cross-reactivity of anti-His antibody (Fig. S17-S20). Therefore, CN:α-6xHis antibody-DyLight-650 complex (2.5 μM, CN:αHisAb650, Abcam ab117504) was prepared by pre-incubating 6x-His-tagged CN with equal concentration of αHisAb650 in CN buffer at ~5 °C for 1 hour. MRBLEs were prepared by transferring a 20 μL aliquot from a 600 μL suspension (~3000-8000 beads depending on pellet size) to a 100 μL PCR strip (20 μL for each CN concentration), exchanging buffer via 3 cycles of iterative pelleting, decanting, and resuspension with 0.1% PBST containing 5% BSA, pH = 7.5 (100 μL), and then left mixing at ~5 °C on a rotator in the same wash buffer overnight (~14 hours). MRBLEs were then buffer exchanged once again with CN binding buffer (50 mM Tris pH = 7.5, 150 mM NaCl, 0.1% TWEEN 20) via 3 iterative cycles of pelleting, decanting, and resuspension.

To perform binding assays, the CN:αHisAb650 complex (at 2.5 μM) was serially diluted into tubes containing BSA-passivated MRBLEs and CN buffer (2 μM, 1μM, 500 nM, 250 nM, 125 nM, 62.5 nM) and incubated for 5 hours on a rotator at ~5 °C. After incubation, MRBLEs were pelleted, decanted, and washed once with 0.1% PBST (100 μL). After a final round of pelleting and decanting, 20 μL of 0.1% PBST was added and the beads were transferred to a quartz microscope slide for imaging. To confirm equilibrium conditions, we measured CN:MRBLE interactions for 6 peptides after incubation times ranging from 30 minutes to 24 hours. Binding appeared to reach equilibrium after ~5 hours (Fig. S6), so all following experiments were performed after an incubation time of 5-6 hours.

### Bead and antibody imaging

MRBLEs were imaged by transferring 20 μL of suspended beads (entire volume) onto a quartz microscope slide (Electron Microscopy Sciences, quartz microscope slide, 75 mm × 25 mm, 1 mm thick, cat. # 72250- 03), placing an additional quartz coverslip (Electron Microscopy Sciences, quartz coverslip, 25 mm × 25 mm, 1 mm thick, cat. # 72256-02) onto the droplet, and then depositing mineral oil around the edges of the coverslip to prevent the sample from drying out during imaging. MRBLEs were imaged largely as described previously^8^ in 11 channels: a bright field channel, a Cy5 fluorescence channel (using a SOLA light engine for excitation), and 9 additional lanthanide channels (435, 474, 536, 546, 572, 620, 630, 650, 780 nm, with exposure times of 500, 1000, 500, 500, 375, 150, 75, 225, 2000 ms, respectively; Semrock, Rochester, NY) using excitation illumination generated by a Xenon arc lamp (Lambda SL, Sutter Instrument, Novato, CA) and directed through a 292/27 nm bandpass filter (Semrock, Rochester, NY) via a UV-liquid light guide (Sutter Instrument, Novato, CA) mounted in place of the condenser using a custom 3D printed holder.

### Image processing (code calling)

Bead images were processed using a custom-built open-source Python software package freely available through PyPI and regularly maintained (Source-code: https://github.com/FordyceLab/MRBLEs). Briefly, MRBLE boundaries were identified from bright field images, followed by quantification of median lanthanide intensities in each channel to identify embedded MRBLE codes. CN binding was then quantified for each MRBLE by calculating the median fluorescence intensity for bound protein associated with the outer shell of the MRBLE.

### Affinity modeling (competitive binding assay)

The affinity (*K_d_*) for each peptide in a MBRLE library was determined via a two-step nonlinear regression using a single-site binding model:

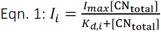

where *I_i_* is the measured fluorescence intensity for each peptide, *K_d,i_* is the dissociation constant for that interaction, [CNtotal] is total CN concentration, and *I_max_* is a global variable representing the fluorescence intensity once MRBLE-pep beads are saturated. First, we determined the global saturation value (*I_max_*), by globally fitting the top 80% highest-intensity MRBLE codes. Next, we globally fit the entire data set using this *I_max_*value to yield an estimated absolute affinity (*K_d_*) for all peptides in the library. Given that the estimated peptide concentration in these assays (~20 nM/x, where x represents the number of species probed) is significantly lower than estimated *K_d_* values of these interactions, we make the approximation [CNtotal]≈[CN_free_]. Differences in binding affinity were calculated relative to a reference peptide using the standard equation (Eqn. 2):

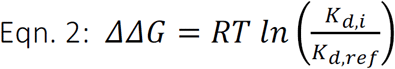

To calibrate measured affinities across multiple assays, we first determined an absolute affinity for a single high-affinity reference peptide (MAGPHPVIVITGPHEE) using the mean of the “triplicate low” value (Fig. 2C), “triplicate high” value (Fig. S3), and value from a binding assay in which this peptide appeared alone (data not shown, *K_d_* = 980 nM). Next, we used this reference value and calculated differences in binding affinity (ΔΔG) to estimate absolute *K_d_* values for each peptide within the calibration library (Table S3-4. Finally, we used calibrated values for several of these peptides (MAGPHPVIVITGPHEE (PVIVIT core and flank libraries), MAGPHPKIVITGPHEE (PKIVIT core and flank libraries), ASGLSPRIEITPSHEL (NFATc2 flank and core library), KRMEPIAIIITDTEIS (AKAP79 flank library and core library) and measured changes in affinity relative to these peptides to estimate absolute ΔΔG values for all peptides in each of these libraries.

### Rosetta-based “Sequence Tolerance” method

For estimating tolerated amino acid substitutions at each position in the two available CN-PxIxIT co-crystal structures with well-defined electron density (positions −1 to 9 of the PxIxIT motif) we used the generalized sequence tolerance module of the Rosetta Backrub server available at https://kortemmeweb.ucsf.edu/backrub^44^. Briefly, the protocol samples amino acid residues in different rotameric conformations on backbone ensembles generated using Rosetta Backrub simulations and records low energy amino acid sequences, from which amino acid frequencies of tolerated substitutions are derived. In contrast to the published protocol, we modified only one position in the peptide at a time, to more closely mimick the experimental measurements. We used ensembles of 100 backbone structures with a kT value of 0.228; self-energies and interchain interaction energies were reweighted using the default scaling factors of 0.4 and 1, respectively.

### Rosetta-based method for estimating the energetic effects of amino acid substitutions (“flex_ddG”)

To estimate changes in binding energy upon mutation (ΔΔG), we used the Rosetta flex_ddG protocol^30^. We systematically substituted all twenty natural amino acids at each position (−1:9) for the two peptides with available crystal structures bound to calcineurin (CN-PVIVIT, CN-IAIIIT), with the remaining positions restricted to the wild-type amino acid residues in each crystal structure. Briefly, the flex_ddG protocol uses the Rosetta Talaris 2014 energy function, minimization with harmonic restraints, Rosetta Backrub simulations to generate conformational ensembles, mutation and optimization of side chain conformations, and another final retrained minimization step. We followed the protocol published in^30^, except using 10,000 Backrub steps instead of the default value of 35000 steps.

### In vivo calcineurin activity assay

PxIxIT peptides were fused to eGFP in pEGFPc1 vector. HEK293T cells were transfected with pEGFPc1 clones, pNFAT-Luc and CMV-Renilla in a 6-well plate format. 18 hours post transfection, 1uM FK506 or DMSO (vehicle) was added to the media as needed. 36 hours post transfection, cells were treated with 1mM Ionomycin and 1mM phorbol 12,13 di-butyrate to activate calcineurin and AP-1 (via PKC) respectively. 6 hours after pathway activation, cells were collected, washed in PBS and re-suspended in DMEM media. 80% of the cells were used to measure luciferase activity and renilla using the Dual-Glo assay system (Promega) with 3 technical replicates. The remaining cells were frozen and stored at −80°C. Cell lysates were prepared in RIPA buffer. 15-20 μg of lysate was analyzed by Western for expression of GFP. GFP signal was normalized to either actin or tubulin. Luciferase activity (normalized to renilla expression) was further normalized to eGFP expression. Data represent at least 3 experimental replicates.

## Acknowledgements

This work was supported by NIH/NIGMS grants DP2GM123641 and R01GM107132 to P.M.F. and R01GM119336 to M.S.C. In addition, P.M.F. is a Chan Zuckerberg Biohub Investigator and acknowledges support from the Beckman Foundation, the Sloan Research Foundation, and P.M.F. and M.S.C acknowledge the support of a joint Bio-X Interdisciplinary Initiatives Fund seed grant. T.K. is a Chan Biohub investigator and supported by NIH/NIGMS grants R01GM117189 and R01 GM110089. We thank Prof. Justin Kinney (Cold Spring Harbor Laboratory) for helpful discussions and software used to generate logos; we thank Prof. Dan Herschlag (Stanford) for essential feedback on the manuscript.

